# Actin protrusions push on E-cadherin to maintain epithelial cell-cell adhesion

**DOI:** 10.1101/644302

**Authors:** John Xiao He Li, Vivian W. Tang, William M. Brieher

## Abstract

Cadherin mediated cell-cell adhesion is actin dependent, but the precise role of actin in maintaining cell-cell adhesion is not fully understood. Actin polymerization-dependent protrusive activity is required to push distally separated cells close enough together to initiate contact. Whether protrusive activity is required to maintain adhesion in confluent sheets of epithelial cells is not known. By electron microscopy as well as live cell imaging, we have identified a population of protruding actin microspikes that operate continuously near apical junctions of polarized MDCK cells. Live imaging shows that microspikes containing E-cadherin extend into gaps between E-cadherin clusters on neighboring cells while reformation of cadherin clusters across the cell-cell boundary triggers microspike withdrawal. We identify Arp2/3, EVL, and CRMP-1 as three actin assembly factors necessary for microspike formation. Depleting these factors from cells using RNAi results in myosin II-dependent unzipping of cadherin adhesive bonds. Therefore, actin polymerization-dependent protrusive activity operates continuously at cadherin cell-cell junctions to keep them shut and to prevent myosin II-dependent contractility from tearing cadherin adhesive contacts apart.

## Introduction

Cadherin family cell-cell adhesion molecules are essential for tissue cohesion and organization throughout life. While the cadherin ectodomain mediates homophilic binding, strong adhesion requires contributions from cytosolic factors including the actin cytoskeleton (1). The apical junction of cell-cell interface is the prominent site of actin polymerization in epithelial cells even long after the junction has been established (2–4). But we still do not fully understand either the architecture or the regulation of the actin at cell-cell adhesion.

Cells generally organize actin into two flavors of architecture: contractile networks that use myosin to generate pulling forces or protrusive networks that use actin polymerization to generate pushing forces (5). Actomyosin contractility, for example, plays a major role in cadherin biology, especially during development when cadherin adhesive junctions propagate tensile forces across interconnected sheets of cells to drive various cell movements (6). Contractility also contributes to junction maturation and stabilization in epithelial sheets (7, 8). Junctional actin polymerization is suggested to build contractile actomyosin (9–14). Observations in cells strongly suggest that cadherin-catenin complexes couple to contractile actin networks and that the complex is under tension (8, 15, 16).

Coupling cadherins to the contractile actin cytoskeleton offers great morphogenetic power to sculp tissues but relying on only contractile forces to stabilize junctions in established epithelia is not fail-safe. In vitro measurements show that piconewton pulling forces stabilize the connection between the cadherin-catenin complex and F-actin (17). In addition, myosin-dependent contractility leads to adherens junction remodeling, perhaps to fortify the junction and make it resilient against tearing (7, 11, 18, 19). But continuing to pull on a broken junction would only tend to propagate the defect which could tear tissues apart (20–23). Cadherin-mediated adhesion is important, and cells tend to evolve safety mechanisms to ensure that important functions remain robust in the face event of a perturbation or occasional failure.

We thus reasoned that actin polymerization-dependent protrusive activity might operate as a safety mechanism to keep lateral membranes of neighboring cells close to each other to promote E-cadherin binding. Protrusions are known to be important for initiating the formation of cell-cell adhesion (24, 25), More recently, protrusive activity was shown to be important for establishing and maintain cell-cell adhesion in endothelial cells (26–28). But epithelial cell-cell adhesion (7, 8, 29–31) is intrinsically more stable than that in endothelial cells (28, 32–34). Whether protrusive activity continues to operate in mature epithelial sheets is not known. Junctional actin assembly in epithelia depends on factors associated protrusive actin networks (2, 3, 23, 35). Thus, we looked for protrusive activities in MDCK (kidney tubular epithelial) cell sheets 3–4 days post confluency with established apical-basal polarity.

## Results

By thin-section electron microscopy we found membrane protrusions of 0.5 µm in length that burrowed into neighboring cells near the apical junctional complex (Fig. 1*A*). Using live cell imaging of F-actin marker UtrCH, we discovered similar filopodia-like microspikes (36) at the apical junctional complex which consists of tight junctions and adherens junctions (Fig. 1*B*) (1). By examining a single labeled cell in a cell sheet, we were able to see membrane structures that are otherwise masked by homogenous labeling like immunofluorescence. Microspikes were detected. Microspikes persist for seconds (mean ± SD lifetime = 10 ± 11 s, *n* = 85) and undergo dynamic elongation, shrinking and pivoting (Fig. 1*C*, Fig. S1*B* and Movie S1). They elongate at the rate of 3.5 µm min^-1^ (Fig. 1*C*). After falling back into the cell body, some microspikes bundle with the junctional actin belt (Fig. S1*B*). Membrane-bound YFP also revealed dynamic protrusions (Fig. 1*D*, Fig. S1*C* and Movie S2). An average cell that is 15 µm wide has ∼50 microspikes (Fig. 1*E*). Blocking actin filament (+) end dynamics with cytochalasin D eliminated microspikes, indicating their dependence on actin assembly (Fig. 1*E*). Simultaneous imaging of actin and E-cadherin showed E-cadherin on the tip of protruding microspikes, indicating that microspikes push on the junction (Fig. 1*C*).

**Fig. 1.**
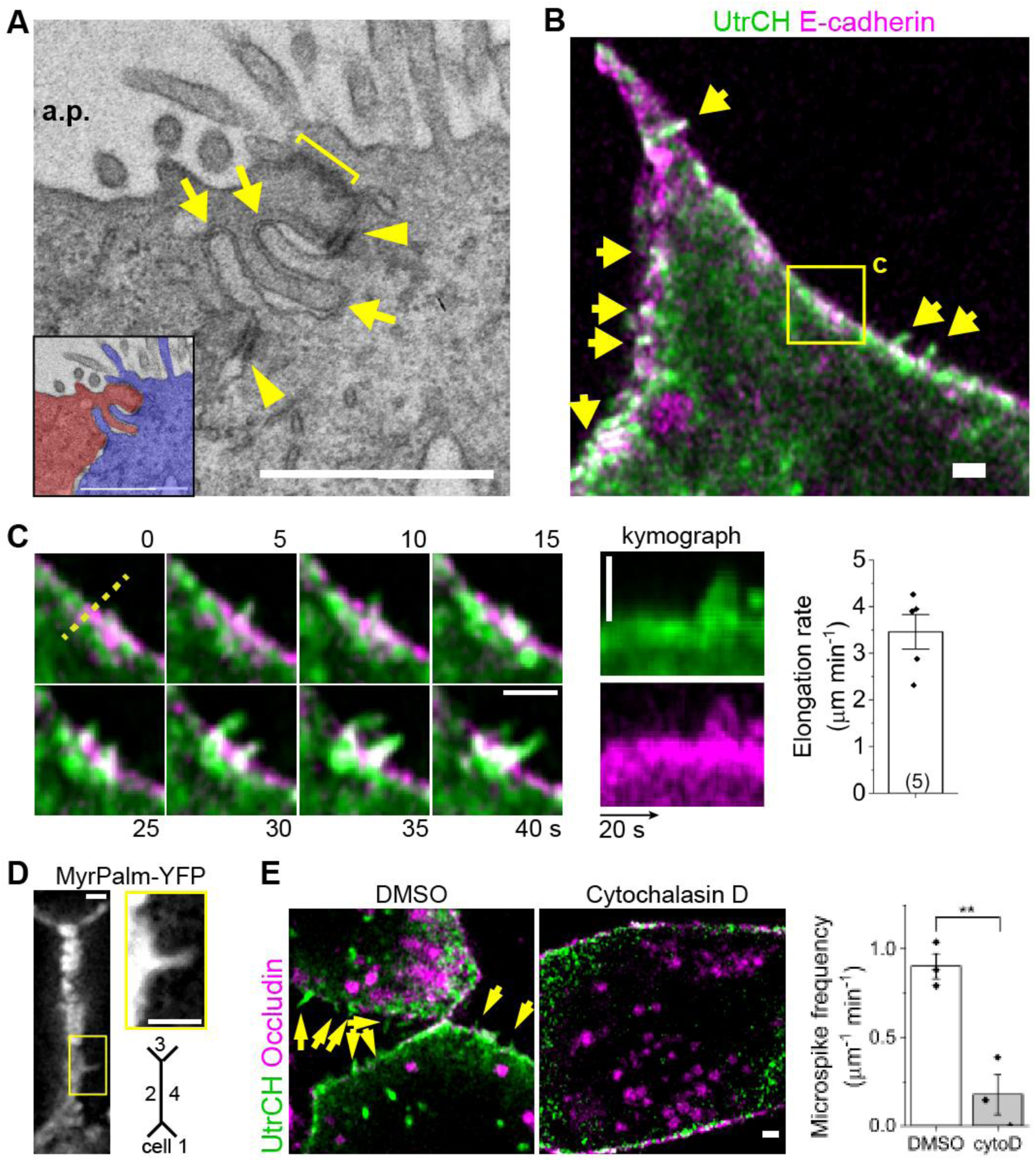
Actin assembly drives microspike protrusions towards apical junction. (*A*) Electron micrograph of confluent MDCK cells (sagittal view). Arrows, protrusions; bracket, apical junction; arrowheads, desmosomes; a.p, apical lumen. (*B*) Microspike protrusions (arrows) in confluent cell sheets sparsely transfected by markers. (*C*) Microspikes elongate and shrink. Kymograph is along the dashed line. Mean ± SEM (*n* = 5 microspikes). (*D*) Membrane-targeted YFP. Inset, membrane protrusion. (*E*) Microspikes (arrows) are reduced by cytochalasin D (*n* = 3 and 3 cells). Mean ± SEM. **P = 0.01, two-sided Mann–Whitney test. Scale bars, 1 μm.

We next asked which actin assembly factors promote microspikes. Previous work identified EVL, CRMP-1 and Arp2/3 as three factors necessary for actin assembly at apical cell-cell junctions (2, 3, 23). Arp2/3 nucleates the formation of new actin filaments (37) while EVL and CRMP-1 form a complex that elongates the (+) ends of existing actin filaments (23). Immunofluorescence showed that all three factors localize to apical junctions (Fig. 2 *A*–*C* and Fig. S2). We therefore tested the role of each factor in microspike formation using RNAi knockdown (Fig. S2*A*). Depletion of EVL, CRMP-1 or the Arp3 subunit of Arp2/3 complex reduced microspikes (Fig. 2 *D* and *E* and Movie S3). Conversely, overexpressing EVL or CRMP-1 led to more microspikes which are longer and more stable (Fig. S2 *D*–*G*). Filopodia and microspikes extend by elongating parallel actin bundles from their (+) ends (36), consistent with EVL and CRMP-1’s function (38). Though Arp2/3 does not make actin bundles, the branched actin it produces serves as the base from which microspikes extend (39, 40).

**Fig. 2.**
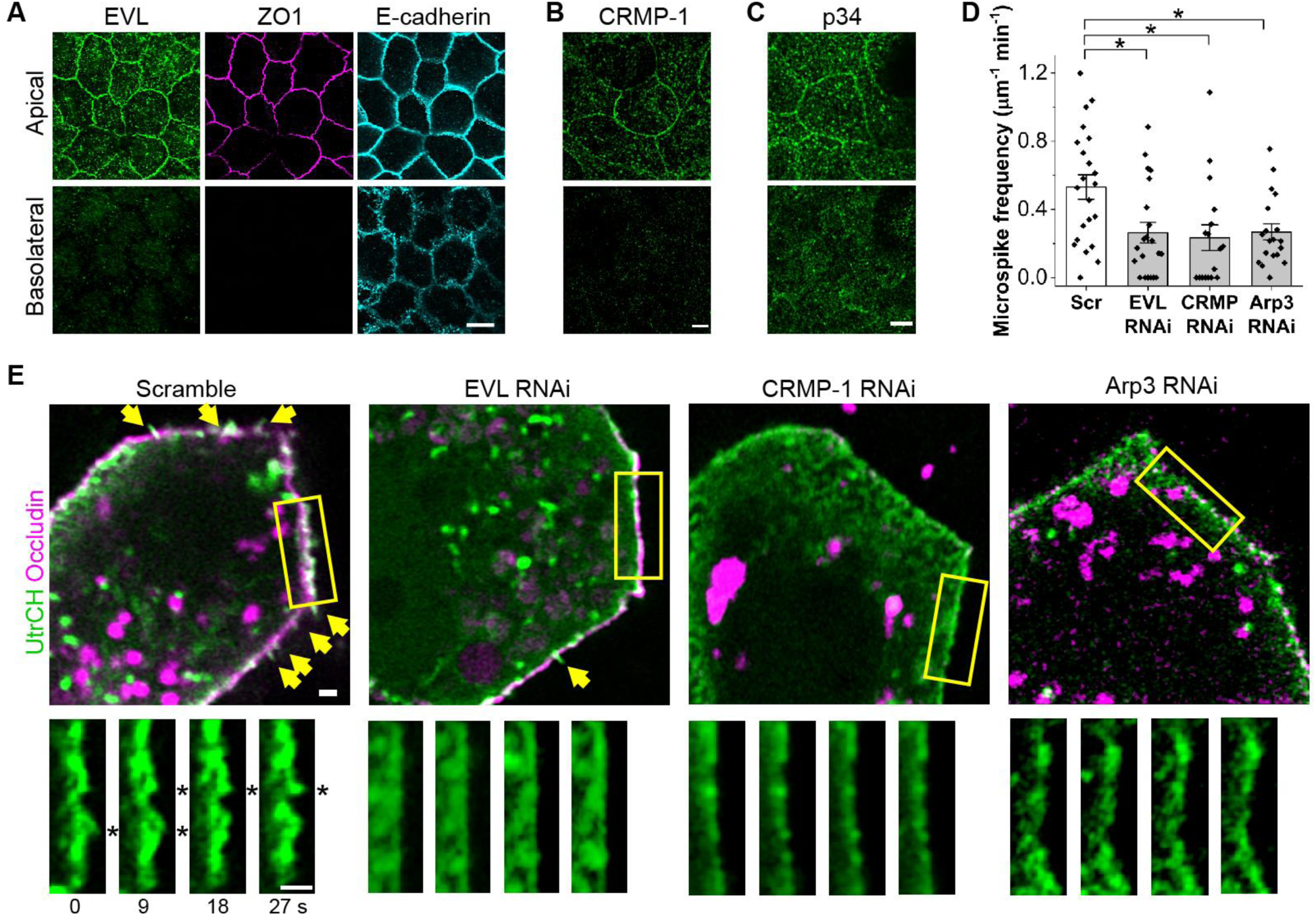
EVL, CRMP-1 and Arp2/3 promote microspikes. (*A–C*) Immunofluorescence of EVL, CRMP-1 or the p34 subunit of Arp2/3 complex with junctional markers. Scale bars 5 μm. (*D–E*) Microspikes (arrows and asterisks) are reduced in cell sheets depleted of EVL, CRMP-1 or the Arp3 subunit of Arp2/3 complex. More EVL- and CRMP-depleted cells have zero microspikes. Mean ± SEM (*n =* 22, 17, 21 and 19 junctions from two experiments). *P = 0.007, 0.02 and 0.04, two-sided Mann–Whitney test. Scale bar, 1 μm.

EVL, CRMP-1 or Arp2/3 depletion also leads to junction failure and cell sheet tearing (21, 23), so we examined the distribution of E-cadherin in these cells. In EVL-depleted cells, cadherin cell-cell contacts are unstable and spontaneously unzip to create deep invaginations in the two cells (Fig. 3 *A* and *B* and Movie S4). EVL-depleted cells have ∼6 invaginations per cell compared to less than 1 in control cells (Fig. 3*B*). Time lapse imaging showed these ∼1 µm deep invaginations (Fig. 3*C*) also occasionally pinching off the plasma membrane to form the large E-cadherin positive intracellular vesicles in EVL-depleted cells (Fig. 3 *D* and *E*, Fig. S3 and Movie S5). These micron-sized vesicles are much larger than clathrin-mediated endocytic vesicles which are 70–200 nm (41). Vesicles extract E-cadherin from the junction in EVL, CRMP-1 or Arp3-depleted cells leading to a weakened junction (Fig. 3*F*).

**Fig. 3.**
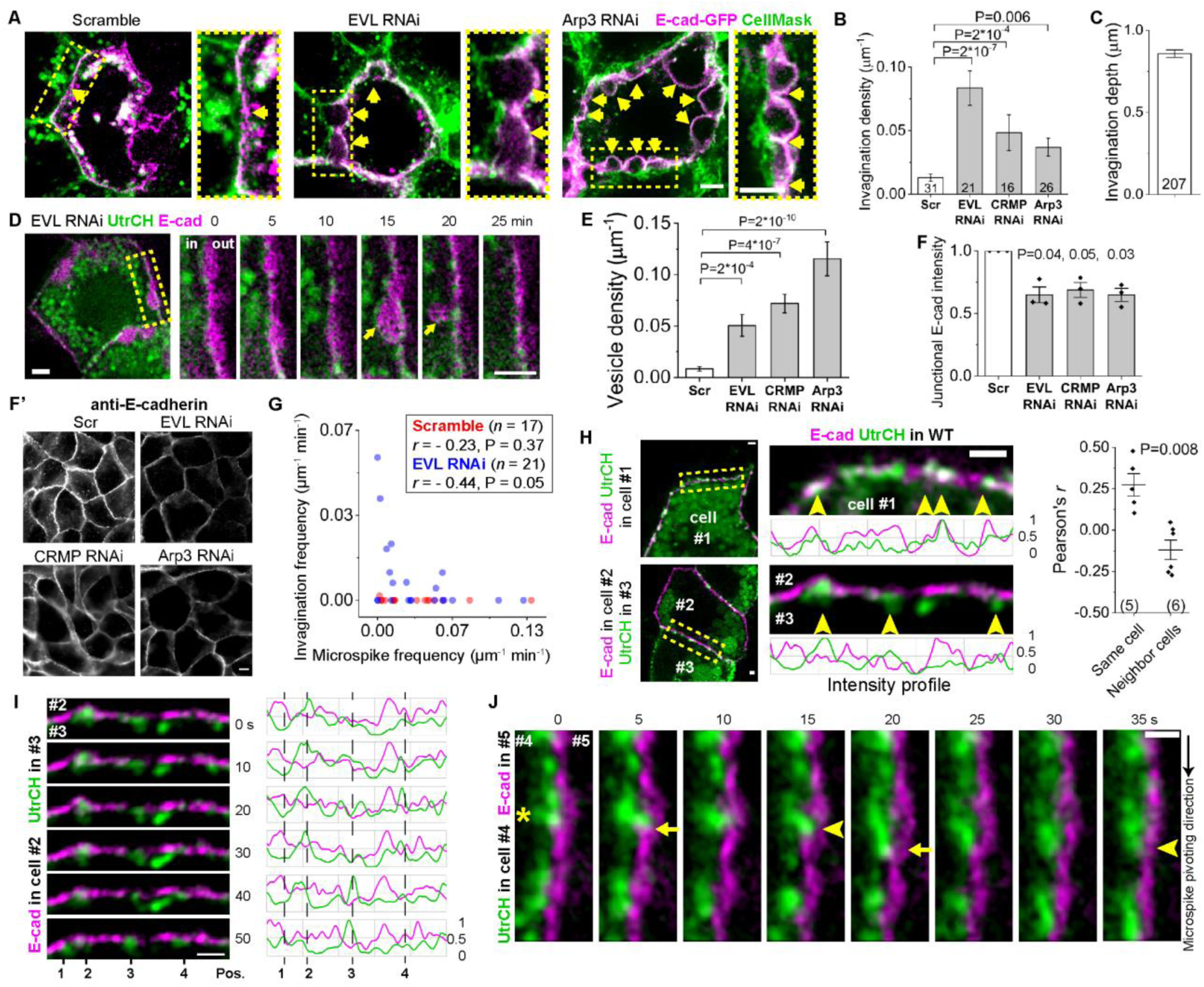
Microspikes push on E-cadherin clusters to repair adhesion failure. (*A–C*) Detached membranes (arrowheads) between E-cadherin-GFP-positive and -negative cells form deep invaginations (*n* = number of cells from two experiments). Scale bars, 2 μm. (*D–E*) The cell-cell junction unzips and internalizes as a vesicle containing E-cadherin (20 minute) in EVL-depleted cells (*n* same as in *B*). Scale bars, 2 μm. (*F–F’*) Junctional E-cadherin immunofluorescence (*n* = 3 experiments, two-sided paired t-test). Scale bars, 5 μm. (*G*) The frequency of invaginations and microspikes in each cell. Pearson’s *r* (*n* = number of cells, P value by two-sided t-test). (*H–I*) E-cadherin and actin markers co-expressed in the same cell (#1) or separately in two neighbor cells (#2 and #3). Microspikes (arrowheads). Pearson’s *r* between E-cadherin and actin intensity profiles along the junction (*n* = 5 and 6 movies). Scale bars, 1 μm. (*K*) E-cadherin and actin markers expressed separately in two cells. An elongating microspike (asterisk) from cell #4 indents the membrane of cell #5 (arrows); indentations rebound (arrowheads). E-cadherin channel shifted by 0.34 µm to the right. Scale bar 1 μm. All bar charts are mean ± SEM with P values from two-sided Mann–Whitney test except for (*F*).

To test whether E-cadherin on the invaginated membrane is no longer in contact with E-cadherin on the neighboring cell, we marked all plasma membrane in the cell sheet with a red fluorescent lipid probe (CellMask) and imaged E-cadherin-GFP transfected individual cells within the sheet. GFP-positive and -negative membranes are from the transfected cell and its non-transfected neighbor, respectively. These two cells’ lateral membranes clearly detached at the invaginations (Fig. 3*A*). Therefore, the deep invaginations are blisters resulting from failures in cadherin-cadherin adhesive bonds.

The number of microspikes is inversely related to the number of invaginations in the cell suggesting that protrusive activity prevents unzipping of cadherin adhesions (Fig. 3*G*). To better understand the relation between microspikes and cell-cell adhesion, we imaged E-cadherin and actin markers expressed in the same cell. The two markers’ intensity profiles along the junction largely overlap indicating E-cadherin present in protruding microspikes (Fig. 3*H*, cell #1 and Movie S6). Furthermore, the changing pattern of intensity profiles supports that E-cadherin is concentrated into dynamic clusters that can appear or disappear over time in an actin-dependent manner (Fig. S4*A*) (12).

Microspikes always correlate with the E-cadherin in the same cell (Fig. 3*H*). However, previous work showed that it is possible for E-cadherin clusters in a *trans* adhesion to dissolve on only one side of the junction (42). Therefore, we wished to know the dynamic relationship between microspikes in one cell and cadherin clusters in a neighboring cell. To examine this, we labeled E-cadherin in one cell and actin in the neighboring cell. In contrast to the coordinated activities between E-cadherin and actin in the same cell, actin intensity in one cell anti-correlates with E-cadherin intensity in the neighboring cell (Fig. 3*H*, cells #2/3 and Movie S6). This suggests that microspikes extend into the gaps between E-cadherin clusters. Their dynamical relation is most easily seen from the intensity profiles which tracks the behavior of four microspikes over time: microspikes at positions 1 and 3 initiated and advanced while microspikes at positions 2 and 4 were extant and subsequently dissolved (Fig. 3*I*). Comparing actin intensity to E-cadherin intensity shows that a microspike initiated within 10 s of the disappearance of E-cadherin (Fig. 3*I*, pos. 1 at 40–50 s and pos. 3 at 10–20 s). In contrast microspikes disappeared within 20 s after filling of the gap in E-cadherin intensity (pos. 2 at 30–50 s and pos. 4 at 20–40 s). The gap is not filled by merging of adjacent E-cadherin clusters as observed when junction is just formed (24) since the clusters remain in their positions (Fig. 3*I*) (12, 43), but rather by E-cadherin projected by microspikes (Fig. 1*C*) (25). Therefore, microspikes are directed to those sites where cadherin-cadherin homophilic binding interactions are absent or failing while restoration of cadherin clusters results in microspike withdrawal (Fig. S4*B* and Movie S7).

We then tested whether microspikes exert pushing forces on the junction. We simultaneously imaged microspikes in one cell and the membrane labeled by E-cadherin in the neighboring cell. Microspikes are associated with the indentations but not the bulges on the neighboring cell’s membrane (Fig. 3*J*) consistent with the electron microscopy (Fig. 1*A*). Furthermore, the indented membrane bounces back once no longer being pushed by the microspike (Fig. 3*J*, 5–15 s, 20–35 s and Movie S8). Therefore, microspikes project cadherins into neighboring cells and indent the neighbor’s membrane in established epithelial sheets, not just in newly forming cell-cell contacts (24, 25).

Unregulated myosin pulling forces can tear epithelial tissues apart (20, 22), we then tested if contractility causes junction unzipping. Inhibiting myosin activity by blocking myosin light chain kinase activity with ML-7 rescued membrane detachment and invaginations in cells that cannot assemble actin protrusions (Fig. 4*A*). Furthermore, myosin II-specific inhibitor blebbistatin functionally rescued junctional E-cadherin in EVL-depleted cells (Fig. 4*B*). Remarkably, blebbistatin also increased junctional E-cadherin in a wildtype background (Fig. 4*C*). This is consistent with the previous work that activating myosin contraction disrupts mature adherens junction (44). Therefore, myosin II-mediated tension weakens E-cadherin homophilic adhesion causing junction unzipping, while in wild type cells these micro adhesion failure events are otherwise repaired by actin protrusions.

**Fig. 4.**
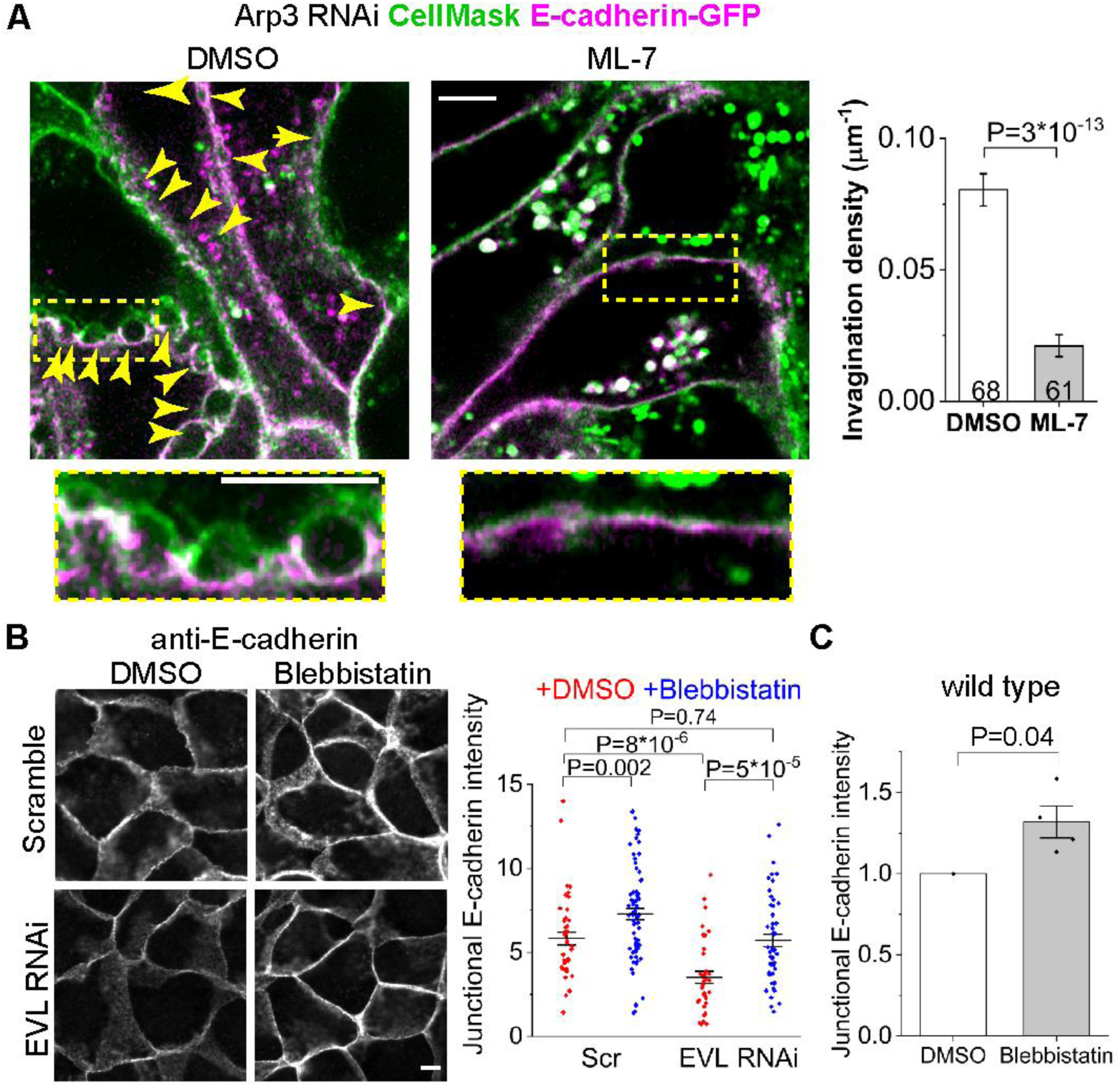
Inhibiting myosin II rescues weakened E-cadherin adhesion. (*A*) Inhibiting myosin light chain kinase rescues detached membranes (arrowheads) between E-cadherin-GFP-positive and -negative cells (*n* = number of cells from two experiments). (*B*) Inhibiting myosin II rescues junctional E-cadherin level (*n =* 44, 75, 36 and 51 junctions). (*C*) E-cadherin immunofluorescence in wild type cells normalized to DMSO (*n* = 4 experiments, with > 30 junctions measured in each experiment; two-sided paired t-test). Scale bars 5 μm. All bar charts are mean ± SEM with P values from two-sided Mann–Whitney test except for (*C*).

## Discussion

We have identified protruding actin microspikes near apical cell-cell junctions and have shown that cadherin adhesive bonds in cells require continuous actin polymerization. We think the protrusive activity serves as a repair mechanism to quickly repair failing adhesive junctions, and we propose a model for how it might work (Fig. 5). Dissolution of a cadherin cluster in one cell triggers the formation of an actin microspike in the neighboring cell. The extending microspike, which contains E-cadherin, extends towards and indents the plasma membrane of the neighboring cell providing an opportunity for re-engagement of cadherin homophilic bonds. Once adhesion is re-established, the microspike withdraws pulling the actin and the cadherins back into the junctional actin belt. This explains why E-cadherin is correlated with microspikes in the same cell but anti-correlated with microspikes in the adjacent cell. In the absence of protrusive activity, myosin II contractile forces overwhelm cadherin adhesive bonds leading to cadherin unzipping. Interestingly then, cadherin homophilic binding in cells does not really require the actin cytoskeleton. Rather, actin polymerization is necessary to constantly push lateral membranes together because contractile forces are always pulling adhesions apart. Once myosin II is inactivated, then protrusive activity is no longer required.

**Fig. 5.**
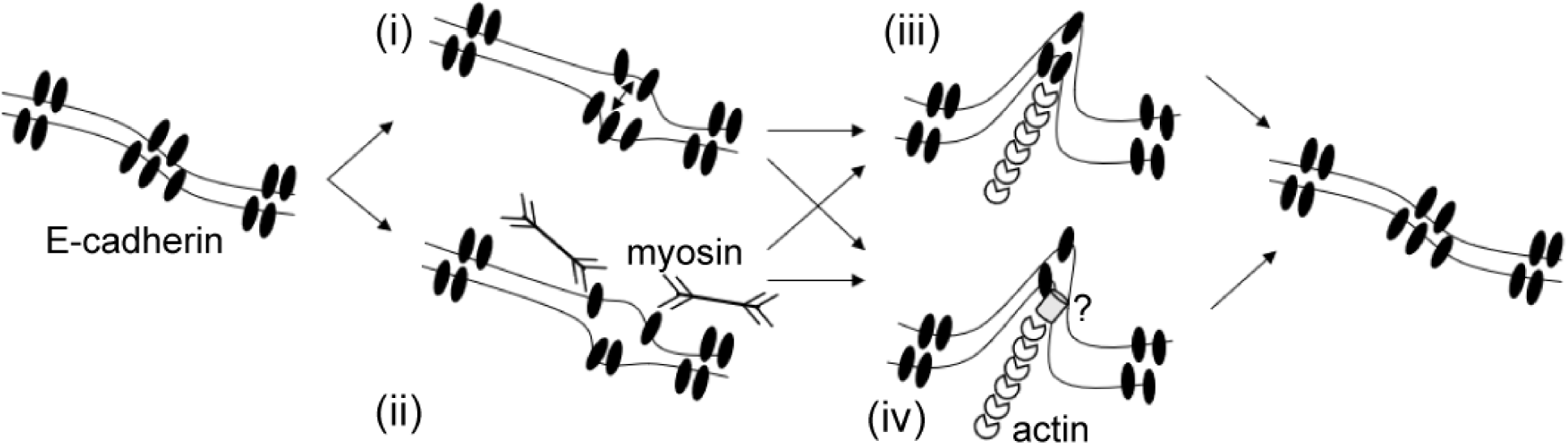
A cell-cell adhesion repair model by actin protrusions. Both (i) spontaneous cadherin adhesive unbinding and (ii) myosin-mediated contractility can cause an adhesion failure. Actin microspikes are triggered by signals mediated by either (iii) unbound cadherins or (iv) unknown factors (question mark). Microspikes push on cadherin clusters into the neighboring cell and promote re-adhesion.

Protrusive activity at cell-cell borders helps explain the otherwise perplexing biochemistry of junctional actin assembly. Contractile actin networks are usually assembled through formin-mediated actin polymerization reactions, and some formins contribute to junctional actin assembly (9, 13, 30, 45). However, other studies also implicated Arp2/3 as a key actin assembly factor at cadherin cell-cell contacts (2, 3, 35). But Arp2/3 is more closely associated with the assembly of protrusive actin networks, not contractile networks (5). Recent work also identified the actin filament elongation factors, EVL and CRMP-1, as important for actin assembly near cadherin cell-cell contacts (23). Again, these factors are most closely identified with protrusive actin (23, 46). Some of the Arp2/3-dependent actin assembly at junctions is linked to membrane trafficking (47). Here we show that these factors continue to drive protrusive activity at cell-cell boundaries long after the cells made contact and long after whole cell motility ceased.

Imaging cadherins and actin in living cells shows that cadherin-mediated adhesion is a far more dynamic and active process than previously understood (12, 42). The physiological necessity of mechanically stable cell-cell contacts motivated searches for molecular mechanisms that would stabilize cadherin mediated adhesion. The cadherin ectodomain alone was sufficient for homophilic binding (48, 49) while experimentally clustering ectodomains in cells could substitute for the cadherin cytoplasmic tail and catenins to mediate strong adhesion (50). The primary function, then, of the actin cytoskeleton was, presumably, to promote cadherin clustering (51) either via a direct linkage to the cadherin-catenin complex (52), or through a corralling mechanism (53). Adhesion was never considered to be static, but the earlier mechanistic work did not predict the fast cadherin-catenin-actin dynamics that operates normally and continuously at cell-cell contacts. A key challenge for the future of adhesion research is to understand the purpose of all these fast actin and cadherin turnover dynamics where actin pulls and pushes on cadherin clusters that are themselves constantly assembling, dissolving, and trafficking to and from the cell surface. Like the actin cytoskeleton itself, cadherin adhesive contacts might be under constant construction. While constant construction requires energy, it would allow cells to rapidly modulate cell-cell adhesion response to signals.

## Methods

### Live cell imaging

Microspikes were imaged at 1-second interval on an LSM 880 laser scanning confocal microscope with an Airyscan detector, a 63×, 1.4 NA oil objective and a heated incubator. Junction invagination and internalization were imaged at 1-minute interval on a V3 OMX microscope with a 100X, 1.4 NA oil objective, two EMCCDs and a heated incubator. Both microscopes allow simultaneous imaging of green and red channels. All cell sheets were grown on Matrigel in glass bottom dishes.

### Microspike labeling and detection

MDCK cells were transiently transfected with UtrCH-GFP to label F-actin while avoiding the probe’s F-actin stabilizing effect. 90% confluent cells were co-transfected with UtrCH and E-cadherin/occludin plasmids and imaged 2–3 days post transfection. Cell contour was segmented by intensity threshold using the actin channel and loaded onto the automatic CellGeo software for microspikes detection (54). Microspikes are protrusions that have length ≥ 0.35 μm, diameter ≤ 0.26 μm, inter-frame displacement < 0.21 μm, tracking gap < 4 s and lifetime > 2 s. The number of microspikes was normalized to junction length and frame interval to give microspike frequency.

### Junction unzipping imaging

Membrane invagination and cytosolic vesicles were imaged in 4-day confluent MDCK cells stably expressing E-cadherin-GFP to reduce cytosolic background. Their numbers were normalized to junction length to give their density.

### Microspike–cadherin correlation

Fluorescence intensity was traced along 5 μm-long junctions to give a pixel intensity profile (*n* = 117 pixels). The intensity profiles of actin and E-cadherin were cross-correlated to calculate their Pearson’s correlation coefficient, *r* = ∑(*I*_1_ - *µ*_1_)(*I*_2_ - *µ*_2_)/*σ*_1_*σ*_2_, where *I, µ* and *σ* are the intensity profile and its mean and standard deviation, respectively.

### Immunofluorescence

EVL, CRMP-1 and Arp2/3 localization were determined by methanol fixation and antibody labeling (23). Apical junctional E-cadherin was fixed with formaldehyde, labeled by antibodies and quantified for its mean intensity on each cell-cell junction.

### Electron microscopy

MDCK cells grown on Transwell were fixed with glutaraldehyde, secondary fixed with osmium tetroxide, en bloc stained with uranyl acetate, embedded and thin-sliced and lastly stained with lead citrate (7, 55).

### Drugs

200 nM cytochalasin D (56), 25 µM ML-7 and 50 µM (-)-blebbistatin were used with 0.1% DMSO as control.

## ACKNOWLEDGMENTS

We thank J. Kemp, A. Nadkarni, K. Prasanth and H. Yu-Kemp for plasmids, cells and antibodies; D. Barry, A. Belmont, K. Cavanaugh, J. Chen, Z. Fu, S. Hilgenfeldt, D. Leckband, T. Saif, W. Sun and D. Tsygankov for discussion; A. Cyphersmith, G. Fried, J. Kim and M. Sivaguru for assistance on imaging; and the National Institutes of Health for funding (R01-GM106106, R01-DK098398).

## Author contributions

J.X.H.L. and W.M.B. designed research; J.X.H.L., V.W.T. and W.M.B. performed research; J.X.H.L. analyzed data; J.X.H.L. and W.M.B wrote the paper.

## Supplementary Information

## Methods

### Plasmids

We inserted shRNAs into pLKO.1 puro vector (RNAi Consortium) against EVL, GCA GGG ATT CAG CCG GAT AAA; CRMP-1, GAT GGA TGA GCT AGG AAT AAA; Arp3, GTA GAT GCC AGA CTG AAA TTA (1) and used a scrambled shRNA as control (Addgene #1864, David Sabatini) (2). We generated SBP-EVL by inserting human EVL (GenBank NM_001330221.1, Origene #RC207094) into the NTAP vector (Agilent #240101) and N-terminal GFP-human CRMP-1 (NM_001313.4) for overexpression (1). beta-actin was cloned into N-terminal GFP pLenti vector (3). The following plasmids are from Addgene: E-cadherin-GFP (#28009, Jennifer Stow) (4); beta-actin-mCherry, mCherry-Occludin-N-10 (#55112, Michael Davison); GFP-UtrCH (#26737), mCherry-UtrCH (#26740, William Bement) (5); LynD3cpV (#37472, Amy Palmer and Roger Tsien) (6, 7); CMV-GFP-NMHC II-A (#11347, Robert Adelstein) (8).

### Drugs

We used 200 nM cytochalasin D for 20 min, 25 µM ML-7 (Cayman #11801) for 30 min and 50 µM (-)-blebbistatin (Cayman #13013) for 1 h with 0.1% DMSO for control.

### Cell culture, plating and transfection

We maintained MDCK II cells in MEM media with L-glutamine (Dr. Sandy McMaster, Cell Media Facility, UIUC) and 5% FBS (Gemini #100-106). The same buffer without Phenol Red is used for live cell imaging. Live cells are in glass bottom dishes (Mattek #P35G-0.170-14-C) and immunofluorescence samples on No.1 glass coverslips. We coated all glass surface with 100 µg ml^-1^ Matrigel (Corning #356237). All dual marker imaging for microspikes was transient expression since UtrCH stable transfection after drug selection induced microspikes in EVL-depleted cells; E-cadherin-GFP for invagination/vesicles and myosin GFP cells were stable expression to reduce cytosolic background. To generate stable cell lines we transfected cells with Lipofectamine 2000 (Life Technologies) and selected with puromycin and/or G418 and collected all surviving cells to make a stable cell line. All knockdown and EVL/CRMP overexpression cells in this article were stable lines. We plated stable cells for immunofluorescence and live imaging at confluency density (4×10^5^ cm^-2^) for four days. To transiently co-express UtrCH and E-cadherin we split cells are split to get 90% confluency the next day. Then we transfected 1 µg of plasmid DNA per marker cells without selection. We imaged cells two-three days post transfection/three days post confluency. The transfection efficiency is low, so we can image single transfected cells in a monolayer with no background. We only imaged cells with medium fluorescence intensity to ensure good signal to noise ratio and avoid artifacts from marker overexpression.

### Immunofluorescence

For EVL/CRMP/p34 localization we fixed/extracted cells in -20° C degree methanol at 2-5 minutes for image clarity (per the protocol of Louise Cramer and Arshad Desai, Timothy Mitchison lab). For all other images we fixed cells with 1% EM grade formaldehyde (Electron Microscopy Services #15710) in pre-warmed Cytoskeleton Buffer (10 mM MES pH 6.1, 138 mM KCl, 3 mM MgCl_2_, 2 mM EGTA) for 15 minutes and then permeabilized and stained cells in 0.1% Triton-X or 0.02% saponin in Wash Buffer (25 mM Tris pH 7.4, 150 mM NaCl) (Cramer and Desai). We mounted fixed sample slides in Prolong Gold antifade (Thermo). We acquired images on a Zeiss LSM 880 scanning confocal microscope with Argon/DPSS/HeNe excitation, an Airyscan detector and a 63×, NA 1.4 objective and automatic settings for Airyscan processing; wide field microscope (AxioImager. M1, ZEISS) with Colibri illumination, a CCD camera (ORCA-ER, Hamamatsu) and a 63×, NA 1.4 objective. We quantified the averaged staining intensity along each junction between two cell vertices. We segmented apical junctions by hand–drawn or ImageJ–generated mask of ZO1 or occludin images. These masks are ∼0.4 µm wide, correspondent to the half width at half maximum of apical junction epifluorescence image. The following antibodies are used: E-cadherin (DECMA-1, sc-59778), Arp3 (sc-10132, Santa Cruz Biotechnology); p34 (#07227, Millipore); AlexaFluor 488/568/647–labeled goat antibodies (Thermo). We raised polyclonal rabbit anti-human EVL or human CRMP-1 full length recombinant proteins^5^ (Pacific Immunology). We produced hybridoma supernatant anti-ZO1 (Dan Goodnough, Developmental Studies Hybridoma Bank #R26.4C) or anti-E-cadherin (Barry Gumbiner, DSHB #rr1) in house (courtesy of James Kemp). We used TRITC-(American Peptide) or Alexa 647-conjugated phalloidin (Cell Signaling Technology #8940).

### Electron microscopy

MDCK cells grown on Transwell-Clear were processed for EM (9, 10). Briefly, transwells were chilled at 4°C for 6 h before fixation (3.75% glutaraldehyde, 150 mM NaCl, and 20 mM Hepes, pH 7.5) at 4°C for 18 h, and then quenched (50 mM glycine and 150 mM Hepes, pH 7.5) on ice for 1 h. Transwells were rinsed in ice-cold distilled water three times, secondary fixed (1% osmium tetroxide and 1.5% potassium ferrocyanide) for 2 h on ice, rinsed four times in ice-cold distilled water, en bloc stained with freshly prepared and filtered 2% uranyl acetate in distilled water on ice for 2 h, and rinsed four times in ice-cold distilled water. Transwells were dehydrated with sequential 5-min incubations in 50, 75, 95, 100, 100, and 100% ethanol at room temperature. Epon-Araldite (EMbed 812) was added to transwells and allowed to polymerize at 60°C for 48 h. Ultrathin sections were cut using a microtome (UltraCut S; Reichert), layered onto carbon-coated copper grids, and stained with freshly made/filtered 2% lead citrate. Images were collected with a microscope (1200EX; JOEL, Ltd.) at 60 kV.

### Live cell imaging

For minute-interval time lapse images we used a wild field OMX microscope with a motorized stage (Applied Precision). For dual labeling we imaged E-cadherin simultaneously UtrCH on two EMCCD cameras, 1 frame per minute for 30 minutes. We deconvolved images with SoftWoRx software (Applied Precision) using the Additive method with automatic iteration settings. We took three z slices 200 nm apart for deconvolution but only used the middle slice for analysis. For all other images we used the same LSM 880 Airyscan confocal microscope for immunofluorescence but with heating at 37° C degree and 5% CO_2_ incubation (Zeiss). For dual labeling we imaged E-cadherin or occludin simultaneously with actin or UtrCH with a dual–pass emission filter, at a frame interval of 1 or 2 s for 60 s. We selected the most apical focal plane with E-cadherin or occludin signal to image apical junctions. Z slices are 190 nm apart.

### Image segmentation for microspike measurement

We pre-processed and segmented cell images of single transfected cells in Fiji for fluorescence correlation analysis (see below). We defined apical junctions as the cell-cell boundary between two vertices. In the time lapse images taken on LSM 880 (z resolution ∼380 nm), only some junctions stayed in focus through 60 s. Thus, we analyzed individual junctions instead of the whole cell and each data point represent a junction. Often only a part of junction was in focus and can be properly segmented, we only used that part. Such part must be > 2 μm to avoid underestimating microspike quantity. Because microspike density is < 1 μm^-1^, so we have sampling frequency > 2. To avoid bias, we only used one part from each junction. We performed walking average of every 2–3 frames and Gaussian smoothing (*σ* = 1 pixel) on movies to reduce noise. We then threshold images with Default or Huang method. Small gaps are the morphologically closed with a circle element of *r* = 2 pixel. We overlay the resulting mask with the raw image to examine its quality. This mask is usually adequate to preserve the fine microspike contour (diameter *∼*0.21 μm); otherwise a unsharp mask filter (*σ* = 15–30 pixel) is applied to the whole dataset to enhance contrast. Within 1 minute, the cell boundary undergoes very small displacement. So that we can register frames with the StackReg plugin (Phillippe Thevenaz) to correct for drift. Registration did not interfere with microspike tracking. Videos 1, 3–5 and 8 are registered with StackReg for presentation. For visualization, we applied Contrast Limited Adaptive Histogram Equalization (CLAHE, Stephen Saalfeld) to enhance microspike contrast in Figs 1*C*, 2*E*, 3*J* and Movies 1, 3 and 7 (block size of 50–90 pixel). We applied unsharp mask to Fig. 1*D* and Movie 2 to enhance contrast.

### Microspike detection

Mask images were automatically analyzed with modified MATLAB– based CellGeo suite to track microspikes (11). Briefly, the BisectoGraph program deduces the cell shape to a tree graph, which transforms the cell boundary into series of convex and concave points. Each convex point is at the end of a branch of the tree, while a joint of branches is the center of a local maximal inscribed circle. Such a circle inscribes the cell boundary at the base of a protrusion and the circle’s radius thus defines the protrusion’s diameter at its base. We then passed the tree graph on to the FiloTrack program to detect microspikes. A protrusion is defined as a microspike if its length > the critical length *L*_*cr*_ and its diameter at the base < the critical radius *R*_*cr*_. Object tracking further enables the identification of same microspike across different frames. Microspike detection threshold is *L*_*cr*_ = 0.35 μm, *R*_*cr*_ = 0.26 μm, inter–frame displacement < 0.21 μm, tracking gap < 4 s, lifetime > 2 s. We measured the frequency, lifetime and length of junctional microspikes from movies of the apical junction using the FiloTrack software. The frequency is the count normalized to junction length and movie duration. The microspike frequency in Fig. 3 is under-sampled than that in Figs 1 and 2, due to different frame intervals (1 s in Figs 1 and 2, 1 minute in Fig. 3 while microspike median lifetime ∼6 s). The estimation of microspike lifetime is affected by the duration of the movie. We used 1-minute long movies but found that a few microspikes persist through the whole movie. 6 s is only of slight under-estimation because 90% microspikes persist < 30 s.

### Junction invagination and internalization

E-cadherin–EGFP labeled membrane invaginations and intracellular vesicles are counted manually from z-stacks. ∼6 slices within 1 micron from the apical junction are regarded as the apical junction. The standard is that deep invaginations must have smooth curvature but not pointed angles and vesicles must have visible hollow cavity. To avoid perturbation of membrane tension, CellMask was not added, thus the invagination density is possibly overestimated (Fig. S3). Their numbers are normalized to the cell perimeter to give their density. The depth of invaginations is the furthest deviation of the invagination from the overall junction contour.

We used double labeling to test whether opposing membranes detach between neighbor cells at the invaginations (Figs 3 and 4). We used CellMask Orange (Thermo #C10045) to label all plasma membrane in the cell sheets and imaged individual cells expressing E-cadherin-GFP within the sheet. Between a GFP-positive cell and a GFP-negative cell, if E-cadherin marker appears on only one of the two CellMask–labeled membranes at the invagination, it shows the membranes have detached.

### Statistical test

Mann–Whitney test was used to compare a sample with control as it does not require assumption of the normality of data (null hypothesis: the medians are equal), except for immunofluorescence or large-sized samples (N > 50, Fig. S*2 C* and *G*). In immunofluorescence due to the difference in absolute intensity values between biological replicates, we normalized the intensity in each experiment to its control (scramble or DMSO) to give relative intensity, *I*, and performed paired t-test (null hypothesis: *I* = 1). For large-sized samples we used the two-sample Kolmogorov test which is more robust (null hypothesis: the medians are equal).

## Movies

### Movie S1

Microspikes elongate/shrink (yellow box) or pivot/fall back into cell body (blue box) at the apical junction. A wild type MDCK cell co-expressing F-actin (UtrCH) and E-cadherin markers in the confluent epithelial cell sheet.

### Movie S2

Membrane protrusions at the apical junction labeled by membrane-targeted YFP (LynD3cpV).

### Movie S3

Apical junction of control (scramble) or cells depleted of EVL, CRMP-1 or the Arp3 subunit of Arp2/3 complex. In cell sheet expressing the shRNAs single cells are labeled for UtrCH and occludin.

### Movie S4

In control cells, E-cadherin tracks UtrCH–labeled protruding microspikes; in EVL–depleted cells, E-cadherin tracks membrane invagination and internalization.

### Movie S5

E-cadherin on membrane invaginations and internalized large intracellular vesicles in EVL– or Arp3–depleted cells.

### Movie S6

UtrCH–labeled microspikes (green) at the junction colocalize with E-cadherin (magenta) in the same cell (upper image) but not with the E-cadherin in the neighboring cell (lower image). In the upper image the cell co-expresses both markers in a cell sheet.; in the lower image the cell at the top is labeled for E-cadherin and the cell at the bottom labeled for UtrCH.

### Movie S7

Cortical actin-GFP accumulates at where junction unzips but subdues after the junction is re-zipped. The gaps in CellMask labeling shows peeling of plasma membranes between the GFP-positive and -negative cells in a cell sheet.

### Movie S8

The cell on the left expresses the UtrCH marker and the cell on the right expresses the E-cadherin marker. The E-cadherin channel was shifted to the right by 0.34 µm. An elongating microspike from the cell on the left pivots clockwise and indents the membrane of the cell on the right; the membrane bounces back after the microspikes pivots away.

**Fig. S1.**
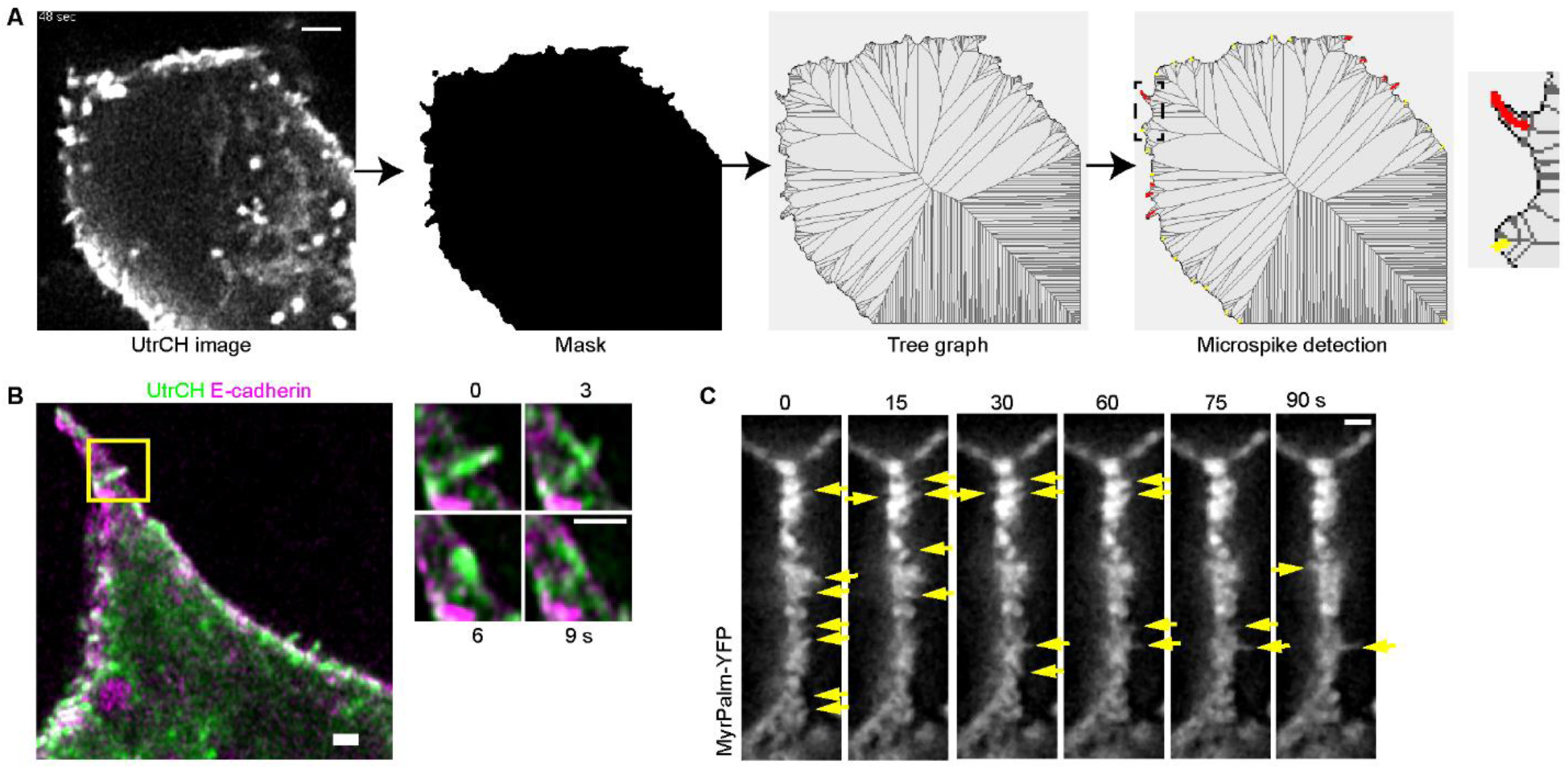
Dynamic microspike protrusions at the apical junction. (*A*) Microspikes are labeled with UtrCH. Image is segmented by thresholding. Cell morphology is deduced to a tree graph with microspikes detected as tree branches meeting the detection standard (red branches). The detection standard includes microspike base diameter and length to exclude branches too short or too wide (yellow branches). (*B*) A microspike falls back into the junctional actin belt. (*C*) Membrane-targeted YFP (LynD3cpV). Arrows, protrusions. Scale bars, 1 μm.

**Fig. S2.**
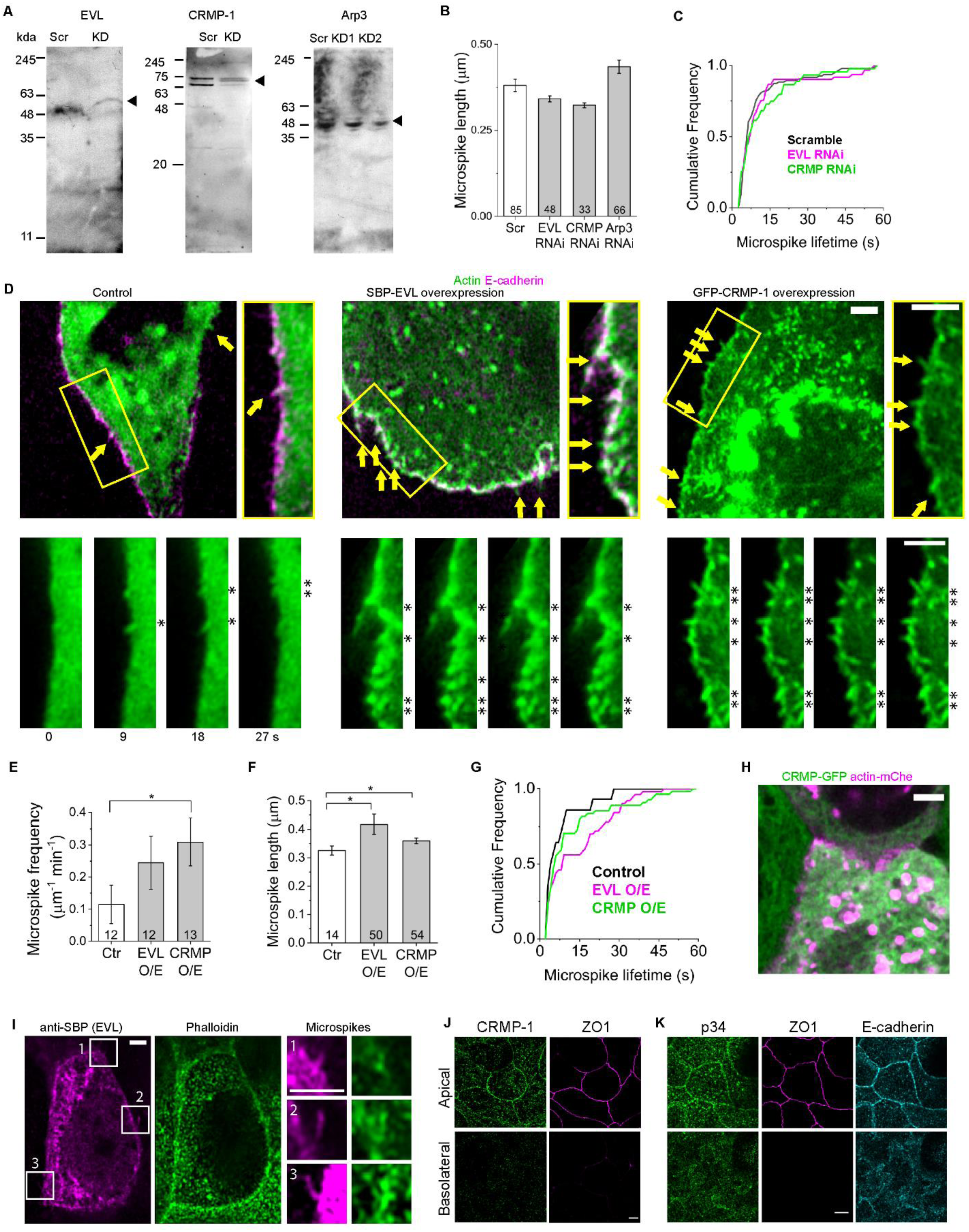
EVL, CRMP-1 and Arp2/3 promote microspikes. (*A*) Whole cell lysate western blot of RNAi knockdown (KD). Scr, scramble. For Arp3, KD2 is the cell line used for all experiments. (*B*) *n* = the numbers of microspikes pooled from two experiments. (*C*) Median ± SD lifetime (s), scramble, 6 ± 11 (*n* = 85 microspikes pooled from two experiments); EVL RNAi, 7 ± 15 (*n* = 48, P = 0.02); CRMP-1 RNAi, 6 ± 12 (*n* = 33, P = 0.06). P value, two-sample Kolmogorov test of cumulative distribution function (bin width = 1 s, *m* = 56 bins). (*D*) Microspikes (arrows and asterisks) in cells overexpressing SBP-EVL or GFP-CRMP-1. Scale bar 2 μm. (*E*) Ctr, parental. *n* = the numbers of junctions. (*F*) *n* = numbers of microspikes. (*G*) Median ± SD lifetime (s), control, 5 ± 7 (*n* = 14 microspikes); EVL O/E, 9 ± 12 (*n* = 50, P = 9*10^-4^); CRMP-1 O/E, 6 ± 14 (*n* = 54, P = 2*10^-7^). P value, two-sample Kolmogorov test of cumulative distribution function (bin width = 1 s, *m* = 59 bins). (*H*) Immunofluorescence of an SBP-EVL overexpression cell. Box, EVL localizes to microspikes. Scale bar 2 μm. (*I*) Live image of GFP-CRMP-1 overexpression cells. Scale bar 2 μm. (*J–K*) Immunofluorescence of CRMP-1 and the p34 subunit of Arp2/3 with junctional markers. Scale bars 5 μm. All bar charts are mean ± SEM. *P < 0.05, two-sided Mann–Whitney test.

**Fig. S3.**
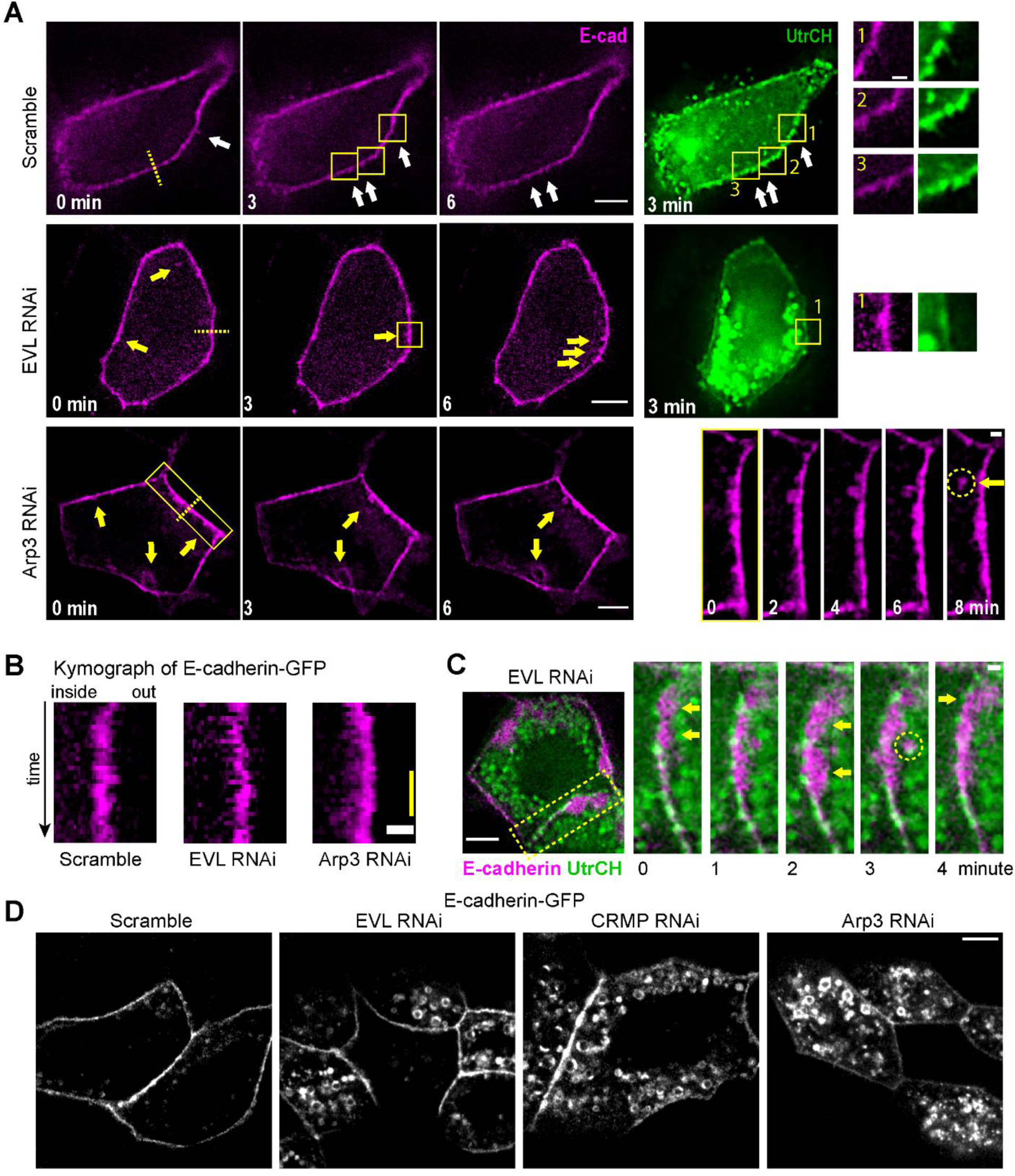
Actin assembly suppresses junction unzipping and E-cadherin internalization. (*A*) E-cadherin tracks microspike protrusions in control (white arrows) and invaginations and vesicles in EVL– or Arp3–depleted cells (yellow arrows). Inset, enlarged boxed regions. Time sequence images of boxed region in the Arp3-depleted cell show an invagination that was internalized (arrow at 8 minute). Scale bars 5 μm except for the enlarged box regions which are 1 μm. (*B*) Kymographs are intensity linescan of the yellow dashed lines, which show the direction of E-cadherin fluctuation. Scale bar 1 μm. Yellow bar 10 minutes. (*C*) In EVL–depleted cells, membranes between neighboring cells detach, invaginate (arrowheads) and internalize as a vesicle containing E-cadherin (circle at 3 minute). Scale bar 5 μm. (*D*) Untreated E-cadherin-GFP cells sheets imaged live showing vesicles. Scale bar 5 μm.

**Fig. S4.**
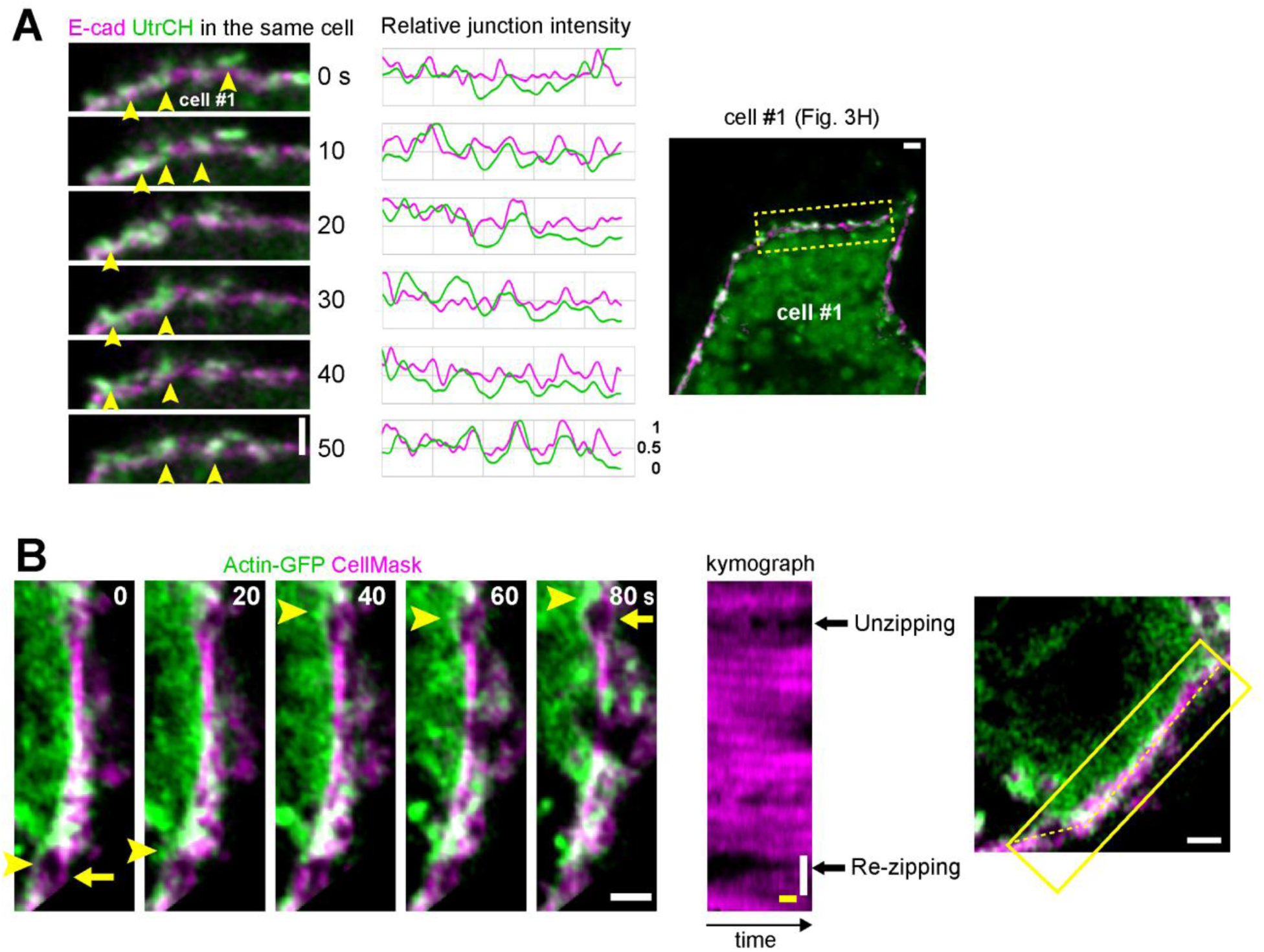
Actin protrusions repair E-cadherin adhesion failure. (*A*) Cell #1 co-expresses E-cadherin and UtrCH. Note that E-cadherin and UtrCH intensities correlate. Arrowheads, microspikes. (*B*) Cortical actin assembles when adhesion fails. In a cell sheet labeled with plasma membrane probe CellMask, actin-GFP labeled single cell formed junction with an unlabeled neighbor. Kymograph is intensity linescan along the dashed line. An unzipping event appears as widening of a gap in the kymograph and a re-zipping events appears as narrowing of another gap. Note that actin accumulates around the unzipping site (arrowhead, 40–60 s) and subdues after the defect is re-zipped (arrowhead, 0–20 s). White scale bars, 1 µm. Yellow temporal scale bars, 20 s.

